# WCRC-25 Is a Novel Luminal Invasive Lobular Carcinoma Cell Line Model

**DOI:** 10.1101/2023.09.15.558023

**Authors:** Ashuvinee Elangovan, Emily A. Bossart, Ahmed Basudan, Nilgun Tasdemir, Osama Shiraz Shah, Kai Ding, Stephen C Purdy, Carolin Meier, Tanya Heim, Carola Neumann, Shireen Attaran, Lauren Brown, Jagmohan Hooda, Lori Miller, Karthik Kota, Tiantong Liu, Shannon L. Puhalla, Grzegorz T Gurda, Lora Heather Rigatti, George Sflomos, Cathrin Brisken, Peter C. Lucas, Priscilla F. McAuliffe, Jennifer M. Atkinson, Adrian V. Lee, Steffi Oesterreich

**Affiliations:** Molecular Genetics and Developmental Biology Graduate Program, University of Pittsburgh, Pittsburgh, PA; Women’s Cancer Research Center, Magee-Womens Research Institute, UPMC Hillman Cancer Center, Pittsburgh, Pennsylvania; Department of Pharmacology and Chemical Biology, University of Pittsburgh, Pittsburgh PA; Integrative Systems Biology Graduate Program, University of Pittsburgh, Pittsburgh, PA; Magee Womens Hospital, UPMC, University of Pittsburgh, Pittsburgh, PA; Department of Pathology, University of Pittsburgh School of Medicine, Pittsburgh, PA; Department of Surgery, Division of Surgical Oncology, Section of Breast Surgery, University of Pittsburgh School of Medicine, Pittsburgh, PA; Swiss Institute for Experimental Cancer Research, School of Life Sciences, Ecole Polytechnique Fédérale de Lausanne (EPFL), Lausanne, Switzerland; Breast Cancer Now Research Centre, Institute of Cancer Research, London, United Kingdom; Fenix Group International, LLC, Philadelphia, PA; Cernostics, Pittsburgh, PA; King Saud University, Riyadh, Saudi Arabia; Takeda Pharmaceuticals, Boston, MA; Centogene, Rostock, Germany; International Association for the Study of Lung Cancer, Denver, CO; Tsinghua University, China; Emplify Health/Gundersen Health System, La Crosse, WI; Mayo Clinic, Rochester, NY; Astellas Pharma, Pittsburgh, PA; Winship Cancer Institute of Emory University, Atlanta, GA

**Author notes:** Corresponding authors, Steffi Oesterreich, PhD Adrian V Lee, PhD, Women’s Cancer Research Center, UPMC Hillman Cancer Center 5051 Centre Avenue, Pittsburgh, PA 15213, USA. Shared first and senior authorships (Ashuvinee Elangovan, Emily A. Bossart, Adrian V. Lee, Steffi Oesterreich).

**Keywords:** lobular breast cancer, pleural effusion, cell line, novel model

## Abstract

Breast cancer is broadly divided histologically into no special type (NST) and special subtypes, of which invasive lobular carcinoma (ILC) is the most common. ILC is distinguished by discohesive, single-file growth from loss of E-cadherin (*CDH1*). Few ILC cell lines exist, limiting mechanistic study of this understudied disease. Here we describe WCRC-25, a novel ILC cell line established from a pleural effusion of a patient with metastatic ILC. While WCRC-25 lacks ER expression, it is molecularly defined as luminal with Her2-like features, and shows anchorage-independent growth and haptotactic migration toward Collagen I. Sequencing identified a *CDH1* Q706* truncating mutation alongside *FOXA1*, *CTCF*, *BRCA2* and *TP53* mutations shared with the patient’s metastases, while copy-number and optical genome mapping revealed structural rearrangements typical of ILC. Transcriptomic AKT-pathway signatures predicted, and cell-based assays confirmed, sensitivity to PI3K/AKT inhibition. WCRC-25 also engrafted and spontaneously metastasized *in vivo*. WCRC-25 is a valuable new tool for dissecting ILC biology and therapeutic vulnerabilities.

## Introduction

About 1 in 8 women in the United States will be diagnosed with breast cancer over their lifetime, with an estimated 43,780 deaths in 2022 (1). Breast cancer is categorized by the molecular and histologic presentation of the tumor. Major histologic subtypes are breast cancer of no special type (NST), also called Invasive Ductal Carcinoma (IDC; ∼80%), and Invasive Lobular Carcinoma (ILC; ∼10-15%), which represents the most common special subtype. ILC tumors are characterized by their growth in a single-file, discohesive manner with stromal infiltration attributed to their hallmark loss of E-cadherin (2–4). Loss of this cell-cell adhesion molecule mostly occurs through genetic deletion, including loss of heterozygosity and truncating or frameshift mutations (5). ILC is more common in older patients, with frequent delayed tumor detection in part due to the unique growth pattern, and with more frequent late recurrences compared with patients with NST (2, 6).

The lower incidence of ILC, relative to NST, previously resulted in this subtype being understudied, but in recent years increased attention has been given to this unique breast cancer subtype. It should be noted that despite being less frequent than NST, ILC affects a large number of women each year (approximately 40,000 annually in the US) and, if considered its own tumor type, would rank as the sixth most common cancer in women (7). Although characterized by favorable prognostic markers, the long-term outcome of patients with ILC is worse than that of patients with NST (8–11). Clinically, patients with ILC are treated in a similar manner to those with NST, and much is still unknown regarding the unique biology and therapeutic vulnerabilities of ILC. Apart from histologic differences, ILC tends to have a lower mitotic index and histologic grade than NST, reflected in overall lower rates of proliferation (12). PTEN loss, activation of growth factor receptor signaling, and PI3K/AKT/mTOR signaling have also been implicated in a significant role in ILC tumor progression (5, 12–17). In addition, our in silico analysis of TCGA and METABRIC data has shown alterations in protein translation and metabolism-related pathways (18).

Although the research community is more motivated than ever to study ILC and significant progress has been made in understanding unique features, further advancement has been hampered by a lack of models that faithfully recapitulate the disease (for a review, see (19)). The Cancer Cell Line Encyclopedia (CCLE) reported 54 NST cell lines and only three ILC cell lines (20, 21), emphasizing the need to expand the inventory of models available to researchers. While there are obvious limitations to using cell lines in cancer research (22), it is important to acknowledge their advantages compared to more complex models, such as their ease of use, low cost, ability to be widely shared with researchers around the globe, enabling independent results validation, and the rapid study of mechanisms underlying phenotypes observed in patients (23). We therefore set out to generate additional ILC cell line models, and here we report the successful establishment and characterization of WCRC-25, a novel ILC cell line generated from a metastatic pleural aspirate from a patient with ILC.

## Materials and methods

### Cell culture

Commercial cell lines were obtained from ATCC (MCF7, MDA-MB-134-VI (MM134), BT474, MCF10-A, MDA-MB-231 (MM231)) and Asterand (SUM44PE). Cell lines were maintained in the following media (Life Technologies) with 10% FBS: MM134 in 1:1 DMEM:L-15, MCF7 and MM231 in DMEM. MCF10-A cells were kept in a 1:1 mixture of DMEM and Ham’s F-12 sans phenol red (Gibco #11039-021) supplemented with 5% Horse Serum (Sigma #H1270), 20 ng/mL EGF (Fisher #EA140), 0.5 mg/mL Hydrocortisone (Sigma #H6909-10ML), 100 ng/mL Cholera Toxin (Sigma Aldrich #C8052-1MG), and 10 μg/mL Insulin (Sigma #I2643-250MG). SUM44PE was maintained as described previously (24)) in DMEM-F12 with 2% charcoal-stripped serum and supplements. Cell lines were routinely tested to be Mycoplasma-free, authenticated by the University of Arizona Genetics Core by short tandem repeat DNA profiling, and kept in continuous culture for <6 months. To overexpress ER, the pCDH-ESR1 lentiviral plasmid was utilized. Puromycin selection was performed until no control cells survived, and selected cells were maintained in medium containing 1.2 µg/mL puromycin, refreshed every two passages. For the Schlegel method, cells were cultured in conditioned media from irradiated J2 fibroblasts combined with F media to generate a complete conditioned medium (25). Murine (BALB/c) stromal-associated fibroblasts (SAFs) stably infected with shScramble or shPRDX1 were cultured in DMEM supplemented with 10% FBS under 5% O2 conditions, and have previously been described (26).

### Acquisition of Patient Samples

Informed patient consent was collected by trained UPMC Hillman Cancer Center/Magee-Womens Research Institute staff. De-identified archival specimens and medical history were obtained, as well as blood and pleural aspirates, from the patient.

### Immunofluorescence (IF) and Immunohistochemistry (IHC)

NST and MCF-10A control cells were plated at 100,000 cells/well on autoclaved glass coverslips (Fisher #12-545-84) in 12-well plates, and ILC cells at 300,000 cells/well. Cells were fixed in 4% PFA for 30 minutes at room temperature before being blocked in blocking buffer (0.3% Triton X-100, 5% BSA, 1X DPBS) for 30 minutes and incubated in primary antibodies for one hour, followed by respective secondary antibodies for 45 minutes at room temperature. Antibody information is provided (Supplemental Table 1). Samples were mounted with DAPI Prolong Diamond (Thermo Fisher #P36962) and confocal images were taken at 40X. Hematoxylin and Eosin (H&E), E-cadherin/p120, and ERα (ER) staining were performed by the University of Pittsburgh’s Pitt Biospecimen Core and UPMC HCC Tissue and Research Pathology Services (TARPS), and reviewed by a certified pathologist (PL).

### Western blot

Proteins were isolated as previously described (16). All samples were quantified using BCA Assay (ThermoFisher #23225) and 25-50 µg per sample were run on 10% homemade SDS-PAGE gels with transfer to PVDF membranes (Millipore #IPFL00010). Membrane blocking was performed with Odyssey PBS blocking buffer (LI-COR #927-40000) for one hour, after which membranes were probed with primary antibodies. This was followed by incubation in 1:10,000 secondary antibodies (Supplemental Table 1) and were imaged on the LI-COR Odyssey CLx Imaging system.

### Phenotypic *in vitro* assays

Soft agar assays were performed in 35 mm culture dishes (Falcon #353001) in a humidified chamber. Enhanced Media (EM, basal media with 20% serum and 2% NEAA) agar at 0.6% final concentration was used for the lower agar layer, and the upper agar layer was generated at 0.4% final concentration in 3% NEAA with 10,000 cells/plate for MCF7 and 50,000 cells/plate for MM134 and WCRC-25. Plates were incubated at 37°C for 21 days, then stained with 0.005% Crystal Violet (Sigma Aldrich #C3886). Dishes were imaged on an SZX16 (Olympus) dissecting microscope with a Nikon DP73 camera under brightfield illumination at 0.8X magnification. Colonies with a radius of 25 µm or greater were counted using cellSens Dimension (Olympus) software, and p-values were calculated with one-way ANOVA.

For Collagen I haptotaxis experiments, the QCM Haptotaxis Cell Migration Assay Collagen I (EMD Millipore #ECM582) kit was used as previously described (24). Cells were plated at a density of 300,000 cells/well in 300 µL of serum-free media; all bottom chambers were also filled with serum-free media. Cells were incubated at 37°C for 72 hours. Excess cells were removed from the top chambers using cotton swabs, and inserts were stained with Crystal Violet (Sigma-Aldrich #C0775) before being imaged on an Olympus SZX16 dissecting microscope and quantified with ImageJ software. Quantifications were normalized to respective BSA controls, and p-values were calculated with one-way ANOVA.

For drug response studies, cells were plated in 50 µL of media at 5,000 cells/well for NST and 15,000 cells/well for ILC cell lines in 2D (Fisher #353072) and Ultra-Low Attachment (ULA) (Corning #3474) 96-well plates. Treatments were added 24 hours post-seeding in an additional 50 µL of respective media. BT474 was included as a control breast cancer cell line with known response to the compounds tested. Alpelisib (Selleckchem #S2814) and AZD5363 (Selleckchem #S8019) were dissolved in DMSO with a final assay DMSO concentration of ≤0.5%. Plates were collected at day 6 and measured by CellTiter-Glo (Promega #PR-G7573). Cell viability values were analyzed following blank cell deductions and normalization to vehicle readings. IC50 values were calculated by nonlinear regression, and statistical differences were evaluated using sum-of-squares global F-test (p<0.05).

### Mutation detection in cfDNA and WCRC-25

Genomic DNA (gDNA) was isolated from WCRC-25 cells using the manufacturer’s protocol for the DNeasy Blood & Tissue Kit (Qiagen #69506) and amplified by PCR using the primer set surrounding exon 13 (Supplemental Table 2). Products were gel-extracted and sent to Genewiz for Sanger sequencing (WCRC-25 Mutant Forward Primer: GAAGCCAAAGATGGCCTTAGA; WCRC-25 Mutant Reverse Primer: GGCATAACTTGGGAGTCTCTTT). The CDH1 Q706* mutation was additionally confirmed by ddPCR, using methods detailed previously (27, 28).

### Mutation Analysis

DNA underwent Ion Torrent sequencing at the University of Pittsburgh Genomics Research Core as previously described (29, 30). Briefly, 20 ng of DNA was used per sample for library preparation using the Ion AmpliSeq library kit (ThermoFisher Scientific) and the MammaSeq primer panel (28, 29). Emulsion polymerase chain reaction (PCR)-based target enrichment was performed using the Ion OneTouch 2 system (ThermoFisher Scientific). Enriched libraries were sequenced using an Ion Torrent Personal Genome Machine (PGM, ThermoFisher Scientific). Raw reads were aligned to the Hg19 reference genome, and variant calling was performed using the Ion Torrent Suite V4.0 proprietary pipeline. Sequencing coverage was analyzed using TarSeqQC v1.18.0 (31, 32). Sequencing depth was aimed at mean coverage greater than 100X. Variant analysis was performed as previously described (29, 30). Briefly, variant call format (VCF) V4.0 files from the Torrent Suite were merged and annotated using CRAVAT (33). Variant filtering was performed to remove pipeline artifacts and common polymorphisms, and to enrich for somatic variants.

Filtered variants were annotated for clinical significance using the PMKB and OncoKB databases (29, 30, 34, 35).

### Copy Number Analysis

Genomic DNA was extracted from WCRC-25 cell pellets using the QIAamp DNA Mini Kit (Qiagen #51306). Briefly, cells were plated in 150 cm^2^ plates. At around 70% confluency, cells were scraped in media and centrifuged at 1400 rpm for 4 minutes, and the supernatant was removed. The pellet was then washed with PBS and centrifuged again before being stored at-80°C. DNA was subsequently extracted using the QIAamp DNA Mini Kit per the manufacturer’s instructions. SNP array analysis was performed at the University of Pittsburgh Genomics Research Core. Briefly, extracted genomic DNA (750 ng) from the WCRC-25 cell line underwent amplification using an Infinium genotyping kit, hybridized to the Infinium CytoSNP-850K v1.2 BeadChip, and analyzed using iScan (Illumina). Raw.idat files were processed using GenomeStudio Version 2.0 (Illumina), using the Hg19 manifest and cluster files provided by Illumina.

CNVpartition v3.20 and CNVRegionReport v2.1.1 plugins were used to generate the final report with logRR values. Segmented data was generated using the copynumber R package (36), and gene-level summary was generated using the CNTools R package (37). The segmentation file from the TCGA BRCA 2015 study (5) (Level3_SNP6_Seg_hg19.txt) was downloaded from FireBrowse (http://firebrowse.org/). Integrative Genomics Viewer (IGV) version 2.11.1 (38) was used for visualization of segmented, genome-wide copy number profiles of the WCRC-25 cell line and TCGA ILC tumors. Copy number alterations (amplifications and deletions) in driver genes in ILC tumors were compared to WCRC-25 as follows: a list of copy-number-altered genes and their frequency in ER+ and ER-ILC samples in the TCGA BRCA 2015 study was acquired from cBioPortal (39). These genes were further filtered to level 4 BRCA driver genes using the OncoVar 2021 database (40).

### Structural Variation Analysis

Optical genome mapping (OGM) was performed at Bionano Genomics Core, San Diego, CA, USA. Briefly, ultra-high-molecular-weight (UHMW) gDNA was extracted from a WCRC-25 frozen cell pellet (1.5 million cells) using the Bionano Prep SP Blood and Cell Culture DNA Isolation Kit (Catalog #80042) as follows: the cell pellet was thawed at 37°C and re-suspended in DNA stabilizing buffer and treated with proteinase K and RNAse A in lysis binding buffer. DNA was bound to a nanobind disk, washed, and eluted in the provided elution buffer (41, 42). UHMW gDNA was labeled using the Bionano Prep DNA Labeling Kit following the DLS protocol. Direct Label Enzyme (DLE-1) and DL-green fluorophores were used to label 750 ng of gDNA. Labeled UHMW gDNA was loaded on a Saphyr chip for linearization and imaging on the Saphyr instrument (Bionano Genomics, San Diego, CA, USA). Structural variant calling was performed using the Rare Variant Pipeline (RVP) with Bionano Solve v3.4 software. Bionano Access v1.7 was used for visualizing structural variations.

### RNA sequencing, sample processing and data analysis

RNA was extracted from FFPE blocks with pathologist-directed microdissection (PL), using the AllPrep DNA/RNA FFPE Kit (Qiagen #80234) according to the manufacturer’s protocol. RNA quality was checked using DV200 metrics performed by the Agilent 4200 TapeStation. At least 25 ng of RNA was used for Exome-capture RNA-sequencing library preparation with Illumina’s TruSeq RNA Access Library kit, and final libraries were sequenced at 40-50 million reads per sample on the Illumina NextSeq 500 platform with a High Output flow cell (2 x 75, stranded, paired-end reads). FASTQ files were mapped and quantified with k-mer-based lightweight-alignment with seqBias and gcBias corrections (Salmon v0.8, quasimapping mode, 31-kmer index built from GRCh38 Ensembl v82 transcriptome annotations). Tximport was used to estimate gene-level expression, and log2-transformed, trimmed-mean-of-M-values-normalized counts per million (log2.TMM.CPM) values were used for subsequent analysis if not otherwise stated. Pearson correlation analysis was performed between each pair of samples using all genes detected. The probability of PAM50 subtype for each sample was called using the genefu package with default settings. Relative strength of cancer hallmark pathways from the MSigDB database was calculated using the GSVA method.

### *In Vivo* Experiments

Mammary fat pad model: 9.4×10^6^ WCRC-25 cells in a 1:1 mixture of PBS and Cultrex were injected into the fourth mammary fat pad on the right side of Female NSG mice (NOD.Cg-Prkdcscid il2rgtm1Wjl/SzJ, cat#:005557 from Jackson Laboratory). Estradiol was supplemented to half of the mice via estradiol pellets (purchased from Innovative Research of America; 17β-Estradiol, 0.5 mg/pellet, 60-day release; cat#: SE-121) inserted into the scruff of mice one week before cancer cell injection. Once tumors were palpable, tumor growth was measured over time via caliper measurements twice/week. Once the average tumor volume for the group reached 1500 mm^3^, mice were sacrificed. Tumors, lungs, livers, and ovaries were collected, fixed in 4% paraformaldehyde, embedded in paraffin, sectioned, and stained with hematoxylin and eosin to assess tumor histology and the presence or absence of metastasis.

For the MIND model, female NSG mice were injected intraductally using a 33-gauge needle with 4×10e5 WCRC-25 cells/mammary gland; no mice died during the course of the experiment. All animal experiments were conducted under the Swiss Institutes of Health Guidelines of Care and Use of Laboratory Animals.

Particular attention was given to preventing animal suffering through daily checks of animals. All mice were maintained and handled according to Swiss guidelines for animal safety, with a 12-h-light-12-h-dark cycle, controlled temperature (22 ± 2 °C), humidity (55 ± 10%), and food and water ad libitum. Experiments were performed under protocol VD3795 approved by the Service de la Consommation et des Affaires Vétérinaires, Canton de Vaud, Switzerland. Tumor growth was assessed by luciferase activity via IVIS imaging. At endpoint, mammary glands were harvested and embedded in paraffin for downstream analysis via H&E and trichrome staining.

### Trichrome Staining

After deparaffinization, the tissue sections were post-fixed in Bouin’s solution for 1 hour at 60°C. Sections were then sequentially stained with Weigert’s hematoxylin to visualize nuclei, Biebrich scarlet–acid fuchsin to stain cytoplasm and muscle, phosphotungstic–phosphomolybdic acid for differentiation and decolorization, and aniline blue to stain collagen. Finally, the sections were dehydrated, cleared, and permanently mounted.

### Data and Cell Line Availability Statement

All data generated in this study are available within the article and its supplementary files, deposited in the Gene Expression Omnibus (GEO) under GSE221512, or available upon request, subject to institutional requirements. The newly generated WCRC-25 ILC cell line is available at [https://www.abmgood.com/Invasive-Lobular-Carcinoma-Cell-Line-WCRC-25-t8074.html].

## Results

### Establishment of ILC cell line WCRC-25

Tumor cells were obtained from a 76-year-old Caucasian woman with Stage IV ILC and metastases to the gastrointestinal tract. The patient was treated with a series of chemotherapies, endocrine therapies, and antibody-based therapies, as depicted in Supplemental Figure S1. The cancer in the breast was bilateral, with one ILC lesion in the right breast (ER+/PR-/Her2-) and a similar lesion in the left breast (ER+/PR-/Her2-), as well as an additional ER-/PR-/Her2-lobular tumor (Figure 1A, Supplemental Fig S2A-C). Lobular histology was confirmed using dual E-cadherin/p120 staining. The disease had also spread to the pleura, which was confirmed to be ILC and showed very weak nuclear ER staining. Cells from the pleural effusion (PE) were used to attempt generation of an ILC cell line (indicated with a “*” in Supplemental Figure S1).

**Figure 1.**
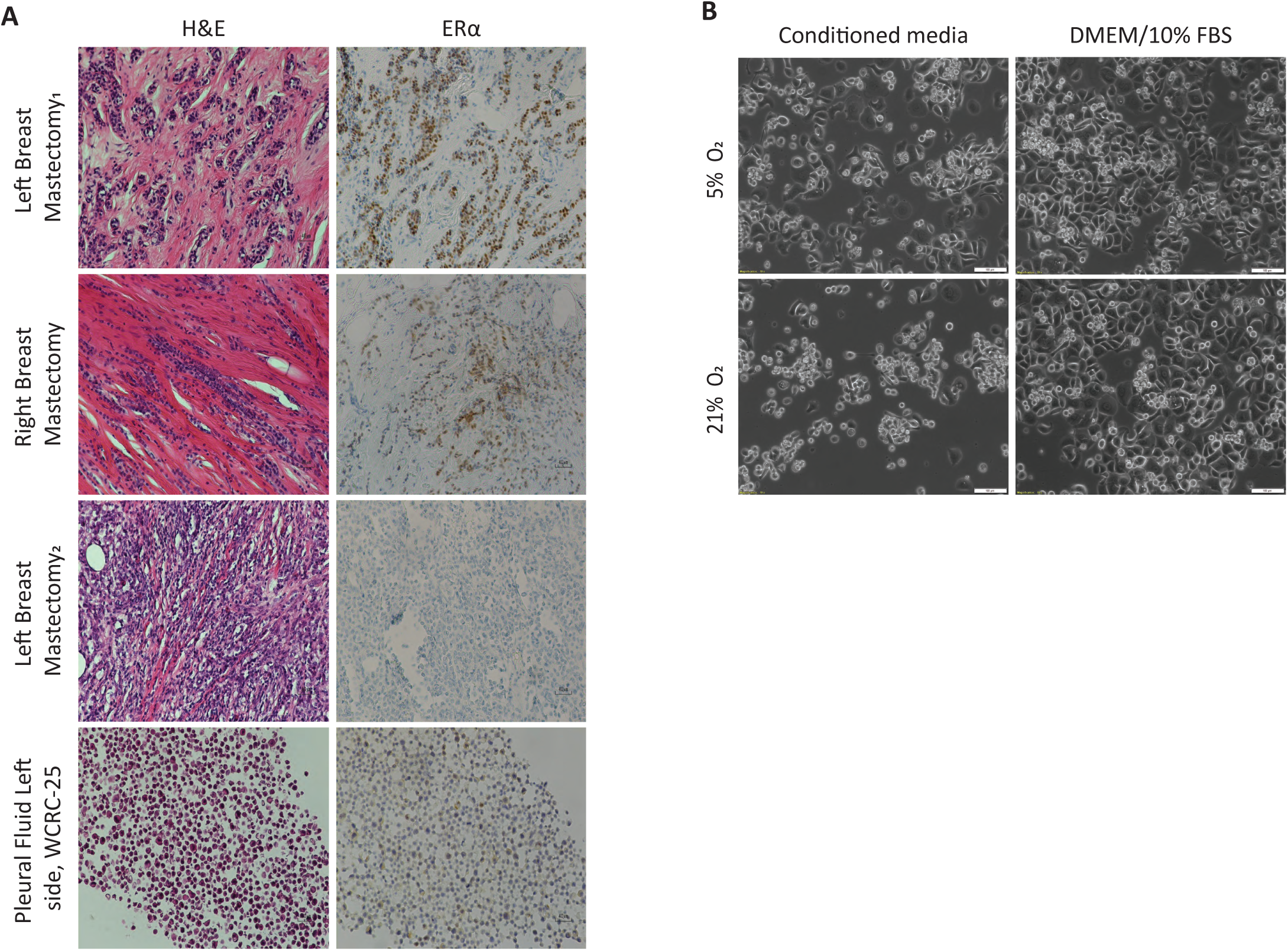
WCRC-25 cell line was established from a metastatic pleural effusion from a patient with ILC. (A) H&E and ERα IHC staining of mastectomy samples and the pleural effusion that led to WCRC-25 cell line establishment (20X magnification). (B) Phase contrast images of WCRC-25 in four growth conditions tested to establish the WCRC-25 culture (10X magnification; bars represent 100 µm). We used hypoxic and normoxic conditions, with 5% and 21% O_2_ (and 5% CO_2_), respectively.

To increase the likelihood of successful generation of a cell line, we cultured the PE cells using multiple different conditions. Parallel cultures were kept in both standard tissue culture (37°C, 5% CO_2_, 21% O_2_, 95% humidity) and hypoxic conditions (37°C, 5% CO_2_, 5% O_2_, 95% humidity) in complete conditioned media (CM) prepared by a modified Schlegel method (43, 44). Cells were plated in standard 2D plates and in ULA plates, with or without supplementation of 10 µM ROCK inhibitor (Y-27632, Enzo Life Sciences #ALX-270-33-M005). We were able to obtain stable lines from the 2D culture in media without ROCK inhibitors grown under hypoxic and normoxic conditions, although cells initially grew slightly better under hypoxic conditions. In order to adjust the cell line to standard cell culture practice, we next cultured these cells in DMEM supplemented with 10% FBS. When comparing cells grown in CM and DMEM/10% FBS, under normoxic and hypoxic conditions, we did not observe major differences in growth and morphology (Figure 1B). As expected, STR analysis reported a unique fingerprint without overlap with existing cell lines (Supplemental Figure S3). We named the cell line WCRC-25, after the Women’s Cancer Research Center at the UPMC Hillman Cancer Center and Magee-Womens Research Institute. The cell line was maintained in DMEM/10% FBS under normoxic conditions, and we proceeded with detailed characterization of this new line.

### WCRC-25 exhibits anchorage independence and haptotactic migration towards Collagen I

Population doubling time, assessed at passages 7, 9 and 11 under the culture conditions described above and across multiple cell densities, was calculated to be approximately 5 days which is comparable to other ILC cell lines (MDA-MB-134: 2 days, SUM44PE: 5-7 days) (Supplemental Figure S4A). Growth was most stable at 10,000 cells/well in a 96-well plate. Based on this experiment, we performed all subsequent phenotypic analyses in normoxic DMEM/10% FBS culture conditions to ensure consistency in cell line comparisons. Late-passage cells showed similar growth rate and cell morphology (Supplemental Figure S4A, B).

We next assessed WCRC-25 growth in 2D and ULA proliferation assays, as existing ILC cell lines have been shown to demonstrate an ability to proliferate well in ULA conditions (24, 45). Compared to MCF-7 and MDA-MB-134 cells, WCRC-25 cells grew slowly in the 2D proliferation assay and were unable to proliferate in ULA (Figure 2A). Although limited growth in soft agar was noted, we observed that WCRC-25 formed more colonies than MCF7 cells, but fewer and smaller colonies than MDA-MB-134 cells (Figure 2B, C). To assess the migration of WCRC-25 towards Collagen I, we performed a haptotaxis assay and observed significant migration (Figure 2D), although smaller than that reported in the ER+ ILC cell line SUM44PE (24).

**Figure 2.**
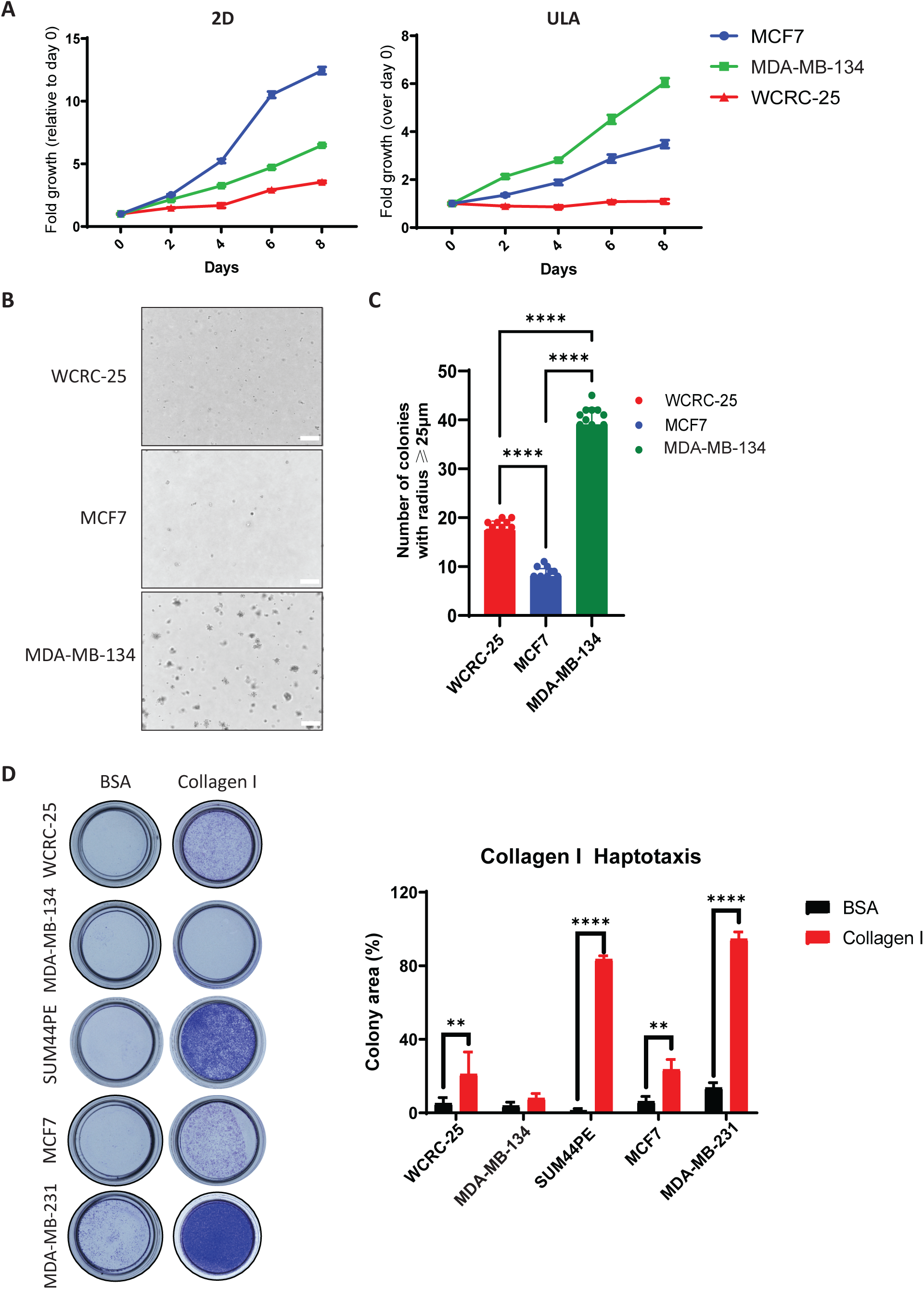
WCRC-25 exhibits anchorage independence and haptotactic migration towards Collagen. **I**(A) MCF7 (5000 cells/well), MM134 (15,000 cells/well) and WCRC-25 (10,000 cells/well) were seeded in 96-well 2D and ULA plates with cell viability assessed on indicated days using CellTiter Glo. Data was normalized to Day 0 quantifications. (B, C) MCF7 (10,000 cells/plate), MM134 (50,000 cells/plate), and WCRC-25 cells (50,000 cells/plate) were seeded in agarose to assess anchorage independent growth. Images were taken on day 21 with quantifications performed for colonies ≥ 25 µm. Representative images are shown and statistical differences evaluated using one-way ANOVA for SD (****p<0.0001, n=3 (each with three biological replicates)). (D) Images and quantification of Crystal Violet-stained Collagen I coated inserts from haptotaxis assays after 72 hours following cell plating at 300,000 cells/well in serum free media. Migrated colony areas were quantified with ImageJ and plotted. Graph shows representative data compared to migration towards BSA from three independent experiments (N=2 biological replicates). p-values shown from Student’s t-test (SEM) was *p ≤ 0.05; **p ≤ 0.01; ***p ≤ 0.001.

This establishment strategy, ie testing multiple culture conditions in parallel, may be broadly useful for deriving additional ILC lines from similarly challenging clinical specimens.

### WCRC-25 is a novel ER-negative epithelial ILC cell line

To assess purity and confirm the epithelial nature of WCRC-25, we stained the cells for a panel of epithelial and stromal markers (Figure 3A). WCRC-25 expressed high levels of the epithelial marker EpCAM and was negative for αSMA, confirming the epithelial nature of the cell line. The cells were strongly positive for luminal markers CK8/18 and negative for the basal marker CK5. Similar to MDA-MB-134, one of the most frequently used ILC cell lines, WCRC-25 also expressed CK14.

**Figure 3.**
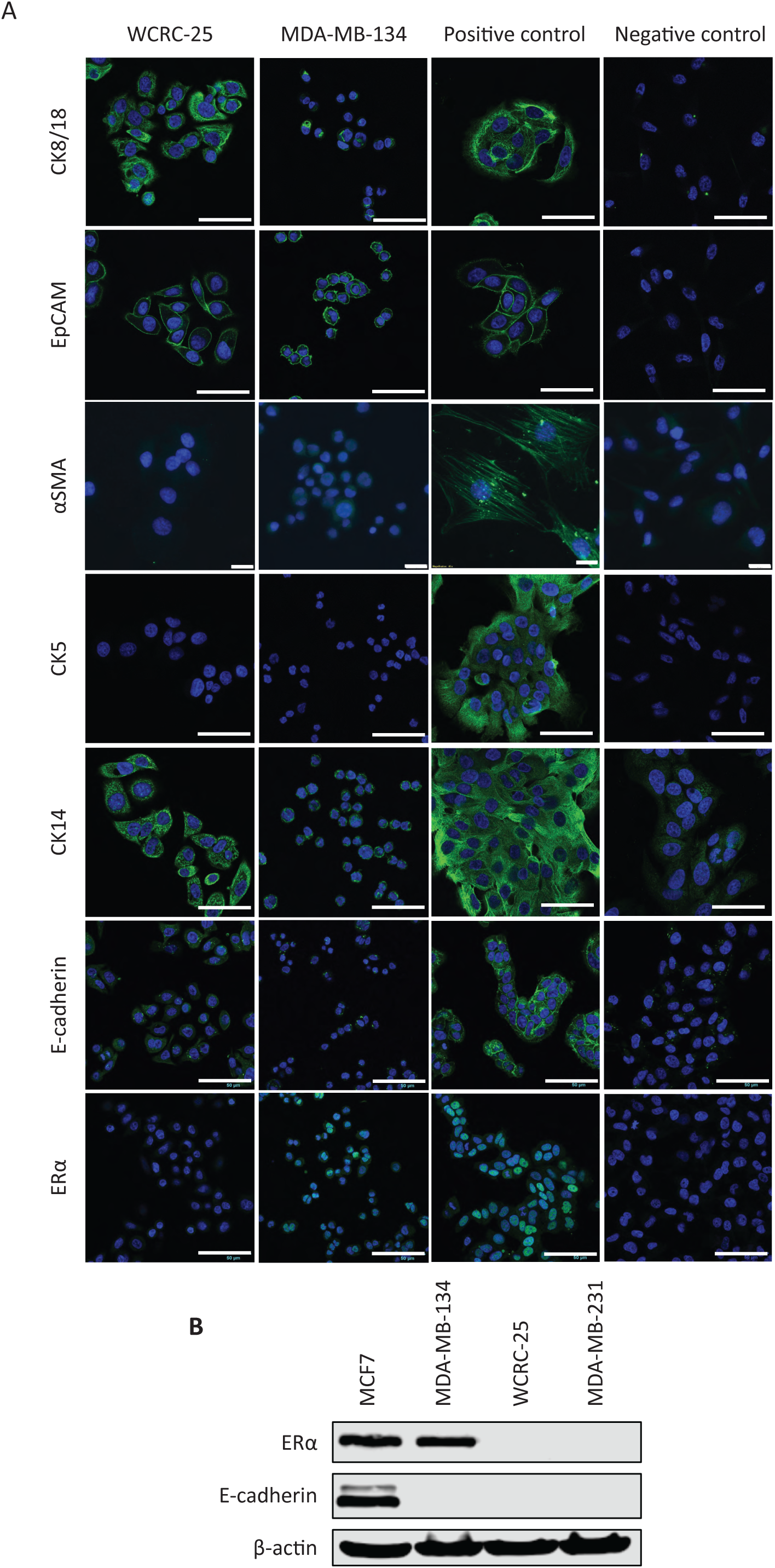
WCRC-25 is an epithelial ILC cell line and lacks ERα expression. A) WCRC-25 cells and appropriate negative and positive controls stained for epithelial markers CK8/18, EpCAM, stromal marker αSMA. CK5, CK14, E-cadherin, and ERα. MM134 cells were included as a classic ILC cell line comparison. Images were taken on a confocal microscope at 40X (CK8/18, EpCAM, CK5, CK14, E-cadherin, ERα) or an inverted fluorescent microscope (αSMA) at 40X magnification. Bars in confocal images represent 100 µm (CK8/18, EpCAM, CK5, CK14) and in inverted images represent 20 µm. Bars in confocal images for E-cadherin and ERα images represent 50µm. WCRC-25 cells and various positive and negative controls were stained for each marker. MDA-MB-231 cells were utilized as a negative control for CK8/18, EpCAM, CK5, E-cadherin, αSMA, and ERα, while MCF7 cells were utilized as a negative control for CK14. MCF-10A cells were used as a positive control for CK5 and CK14 while shPRDX1 SAFs/CAFs were used as a positive control for αSMA. For CK8/18, EpCAM, E-cadherin and ERα, MCF7 cells were utilized as a positive control. Representative images are shown of n=2 independent experiments. (B) Immunoblot for ERα, E-cadherin and β-actin with WCRC-25, MCF7, MM134 and MM231.

We next assessed expression of E-cadherin and ER in WCRC-25 (Figure 3A, B). In contrast to MCF-7, which showed the expected strong membranous staining of E-cadherin, we only observed very weak, punctate staining in the cytoplasm in WCRC-25, similar to that observed in MDA-MB-134. Absence of full-length E-cadherin protein was also confirmed by Western blot in WCRC-25 (Figure 3B). ERα expression was lacking in WCRC-25 as shown by immunofluorescence (Figure 3A) and immunoblotting (Figure 3B). Since there was low ER expression in the PE from which WCRC-25 was generated, we attempted overexpression of ER in WCRC-25. Constitutive expression of HA-tagged ER by lentiviral transduction was initially successful in WCRC-25, but the expression was lost over time despite constant selection pressure (Supplemental Figure S5A). An ERE-TK assay showed no effect of E2 or ER antagonists on ER activation (Supplemental Figure S5B), confirming the absence of functional ER in WCRC-25. Altogether, we confirmed the epithelial nature of WCRC-25, a novel ER-negative lobular cell line.

These data establish WCRC-25 as a bona fide epithelial, ER-negative ILC model, setting up the genomic as well as functional analyses of ER-independent vulnerabilities described below.

### WCRC-25 carries a CDH1 mutation in addition to other DNA structural variations

Next, we analyzed loss of E-cadherin (*CDH1*) - the hallmark of ILC - in more detail in WCRC-25. We observed complete loss of E-cadherin expression (Figure 3B), suggesting that WCRC-25 harbors *CDH1* loss of heterozygosity and a point mutation, as commonly seen in other ILC cell lines (46–48). Sanger sequencing confirmed a *CDH1* nonsense mutation at Q706 with a C>T transition leading to the formation of a premature termination codon (PTC) (NM_004360.4:c.2240C>T, p.Q706*) (Figure 4A). Using ddPCR technology, we were able to detect the CDH1 Q706* mutation in the cell line, with a 97.9% mutant allele frequency (AF) (Figure 4B).

**Figure 4.**
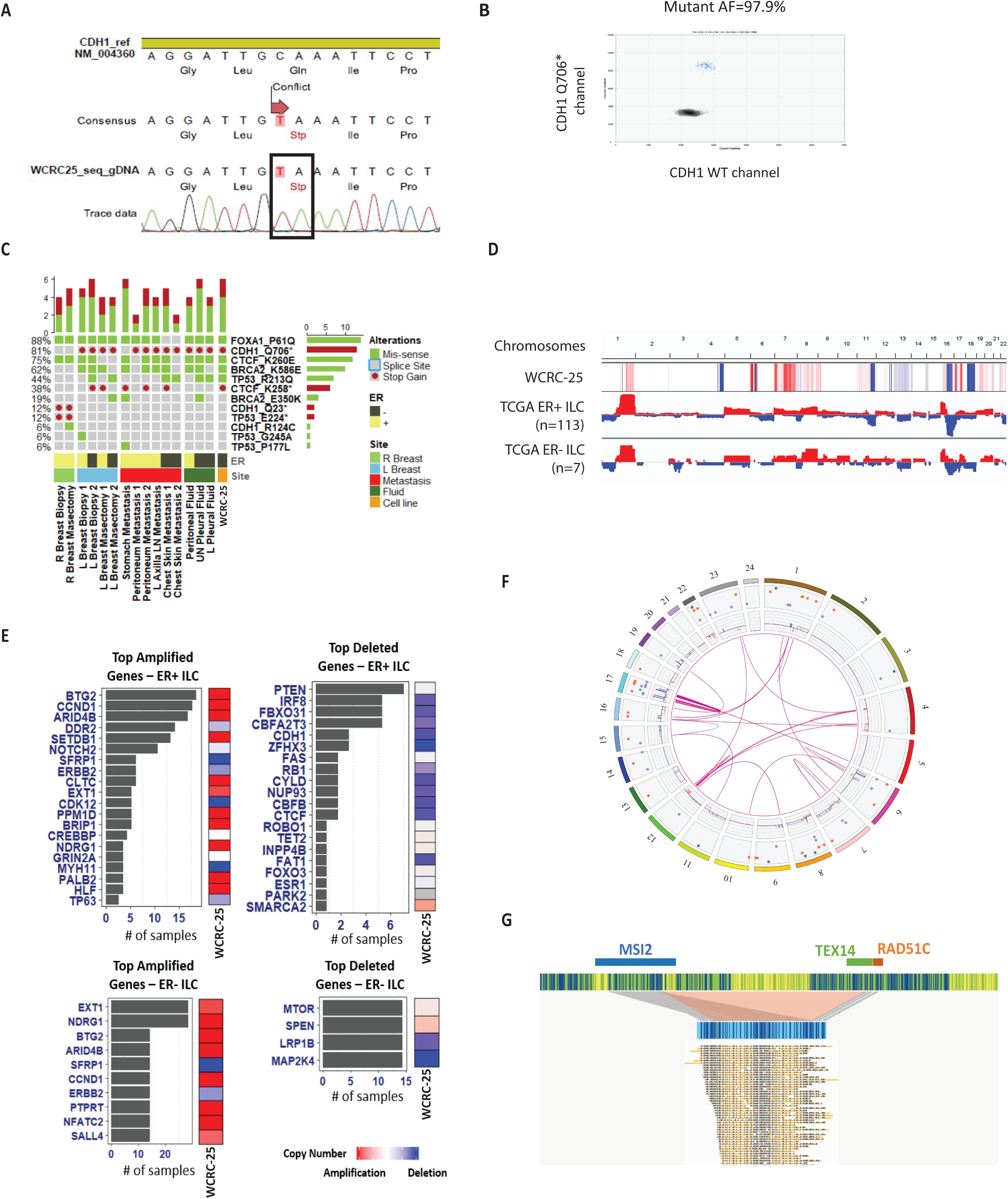
A novel *CDH1* mutation at Q706 is detectable in patient cfDNA and in the WCRC-25 cell line. (A) Detection of *CDH1* Q706* mutation in WCRC-25 gDNA by Sanger sequencing, denoted in chromatograms with a black box. (B) ddPCR performed in the WCRC-25 cell line showing a 97.9% CDH1 Q706* mutant allele fraction. (C) Oncoprint showing the top 5 DNA alterations identified using MammaSeq. Each row represents a unique alteration (shown by gene name and the associated protein change) and each column represents a sample. Samples are ordered as those from right breast, left breast, metastasis, fluid and cell line. (D) Genome-wide copy number profiles of WCRC-25 cell line compared to TCGA ER+ and ER-ILC tumors. Red represents copy number gain and blue represents copy number loss. (E) Barplots representing alteration (amplification or deletion) frequency (# of samples with alteration) of driver genes in TCGA ER+ and ER-ILC samples. Heatmaps adjacent to barplots represent copy number of corresponding genes in the WCRC-25 cell line. Red represents amplification and blue represents deletion. (F) Genome-wide structural variation (SV) displayed using the Circos plot. (G) Detailed description of the putative fusion identified in chromosome 17 of WCRC-25 via Bionano optical gene mapping.

To identify additional mutations in clinical samples from the patient from whom WCRC-25 was derived, and to determine how well the WCRC-25 model reflected the genomics of the patient’s tumor, we performed targeted DNA sequencing using the MammaSeq panel (29, 30). We isolated sufficient DNA from breast biopsies and the mastectomy sample, from the tumor in the lymph node, and from metastases in peritoneum, stomach, and skin. All samples passed quality control assessment (average coverage > 100X). Mutations were annotated using CRAVAT (33) and enriched for somatic variants. A total of 61 unique mutations in 32 genes were identified across patient specimens (Supplemental Figure S6A, top 30 most frequently mutated genes, samples are ordered as those from the right breast, left breast, metastasis, fluids, and cell line). Unique mutations in the top 5 most frequently mutated genes (*CTCF, CDH1, FOXA1, TP53*, and *BRCA2*) are shown in Figure 4C and included mutations in *CDH1* and *FOXA1*, which have previously been associated with ILC (5). Interestingly, we identified two unique *CDH1* mutations: *CDH1*-Q23*, seen only in the right breast biopsy and mastectomy samples, and *CDH1*-Q706*, seen in the left breast biopsy and mastectomy as well as in all metastases (with the exception of stomach), fluids and the WCRC-25 cell line, as previously observed via ddPCR. The most frequently identified mutations, including *FOXA1* P61Q, *CTCF* K260E, *BRCA2* K586E, and *TP53* R213Q, were maintained in WCRC-25.

Given that copy number variations are important drivers of breast cancer (49, 50), we performed the Illumina 850K cytoSNP analysis to better understand the copy number landscape of the WCRC-25 cell line. A genome-wide copy number profile of the WCRC-25 cell line as compared to TCGA ILC tumors (Figure 4D) shows that overall, WCRC-25 captured many of the gains and losses observed in ER+ and ER-ILC, including gains in 1q, 7p, 7q, 8q, 11p, 16p, 19p, 19q, 20p and 20q, and losses in 4q, 11q, 13p, 13q, 16q, 18p and 18q. Breast cancer driver gene copy number alterations were also identified in the WCRC-25 cell line. A list of frequently altered genes in TCGA ILC tumors (5) was acquired from cBioPortal and filtered for pathogenic driver genes using the OncoVar database (40). We observed that WCRC-25 captured the respective alterations (amplifications or deletions) in the top copy number altered driver genes in ILC tumors as shown in Figure 4E. The top 5 conserved amplified and deleted genes included *BTG2, CCND1, ARID4B, SETDB1* and *CLTC*, and *PTEN, IRF8, FBXO31, CBFA2T3* and *CDH1*, respectively.

To gain further insight into DNA structural variations (SV), we performed Bionano optical mapping of DNA from the WCRC-25 cell line. This analysis identified a total of 154 SVs (Figure 4F), with the most common alterations being deletions and duplications, followed by translocations (Supplemental Figure 6B). We also observed many intra-chromosomal translocations and identified many putative fusions in chromosome 17 of WCRC-25, suggesting a chromothripsis-like event (Figure 4G, Supplemental Figure S6C). An MSI2-TEX14 putative fusion identified in chromosome 17 is shown in Figure 4G. This fusion resulted from a deletion spanning from Chr17: 55655097 – 56731701. Given MSI2’s previously described role in altering ER activity (51), this fusion gene could have played a functional role in the development or progression of the tumor.

Collectively, the *CDH1* truncating mutation and the ILC-typical copy number landscape identified here confirm that WCRC-25 recapitulates the genomic hallmarks of ILC while also harboring previously undescribed alterations warranting further study.

### RNA sequencing reveals enhanced AKT signaling that predicts, and is functionally confirmed, by susceptibility of WCRC-25 to PI3K/AKT inhibition

To investigate how closely RNA expression in WCRC-25 would resemble the transcriptomic features of the patient’s tumor, Pearson correlation analysis was performed on RNA sequencing data from the WCRC-25 cell line and the patient’s specimens, including the primary breast tumor, peritoneal metastases and fluid, chest skin metastases, lymph node metastases, stomach metastases and PE. Gene expression of WCRC-25 was most similar to the gene expression profile of the PE (Figure 5A), in line with it being the source of the cell line. PAM50 subtype assignment showed that all primary breast tumors were of luminal A or luminal B subtype (Figure 5B), while some metastatic samples showed subtype switching. The majority of the metastases maintained luminal features; however, one chest skin metastasis switched to the basal subtype, and another chest skin metastasis, a stomach metastasis and left pleural fluid gained features from the HER2 subtype. Of note, both the WCRC-25 cell line and the PE had luminal features (LumB) shared with the primary tumors, but had also gained HER2-like subtype features.

**Figure 5.**
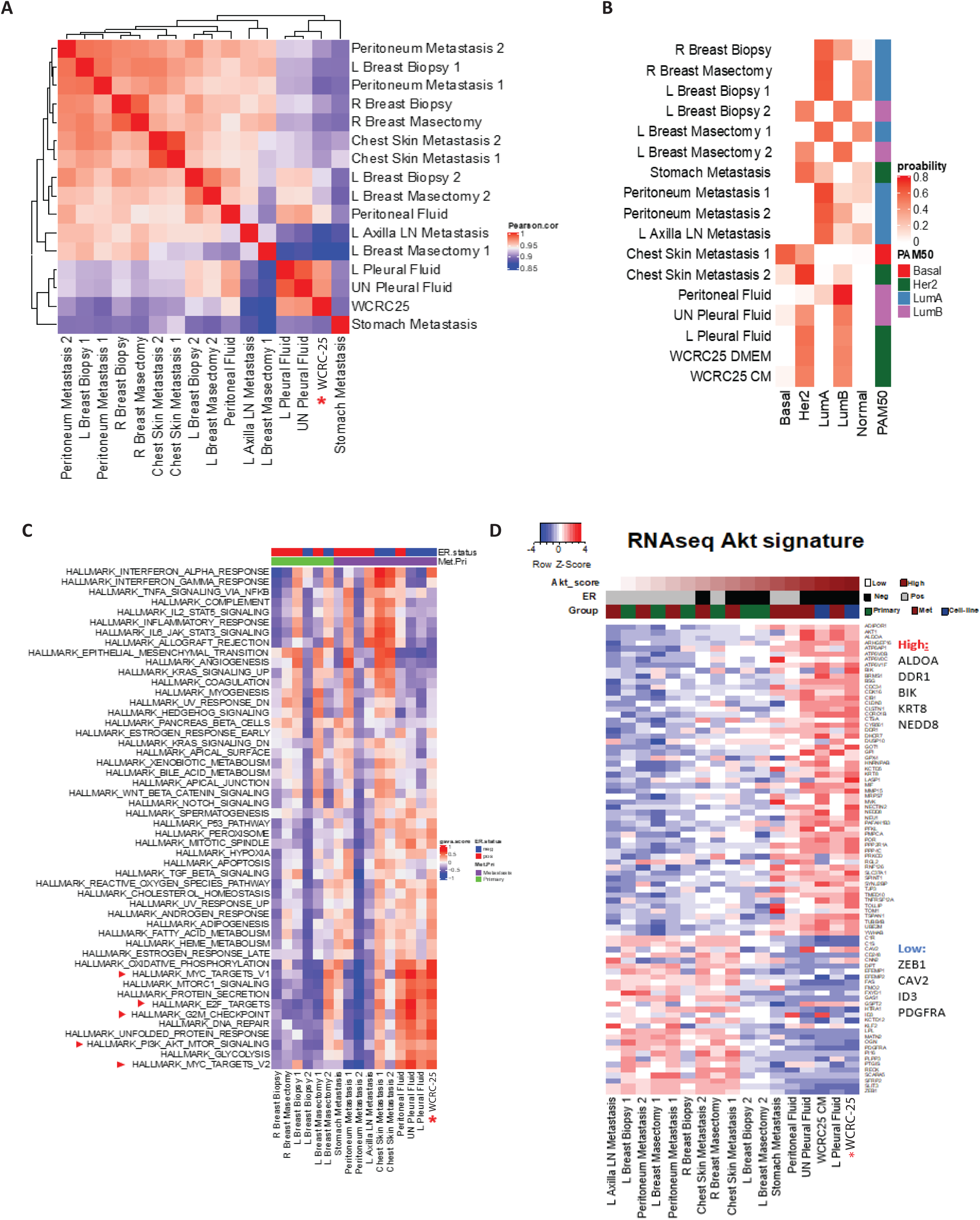
RNA sequencing reveals enhanced AKT signaling and intra-lesion heterogeneity in clinical specimens. (A) Pearson gene expression correlation matrix between each sample pair. Hierarchical clustering was performed based on the correlation matrix. (B) Relative probability of PAM50 subtypes across samples, with the highest probability for each sample noted to the right of the figure. (C) GSVA score of cancer hallmark pathways across samples, with primary tumors and metastases grouped together. Pathways related to cell proliferation are highlighted with red arrow. (D) Z score of genes upregulated and downregulated by AKT across samples. Columns were ordered based on their relative AKT score.

To examine the relative strength of cancer-related pathways across primary tumor and metastases, and to compare this to what is seen in WCRC-25, GSVA scores of cancer hallmark pathways were calculated and compared across samples (Figure 5C). Metastases generally showed higher activity of pathways related to cell proliferation, including MYC targets, E2F targets, G2M checkpoints, and PI3K-AKT-mTOR pathways. Hyperactivation of the PI3K-AKT pathway is a common mechanism driving progression and treatment resistance of breast cancer metastases. Genes upregulated by AKT (52) showed consistently high expression in samples with high AKT pathway activity, especially ALDOA, DDR1, BIK, KRT8, and NEDD8, while the opposite was true for genes downregulated by AKT (52), with ZEB1, CAV2, ID3, and PDGFRA being the most downregulated genes (Figure 5D).

Given this activation of the AKT signature, we directly tested its functional relevance by assessing cell viability with pathway inhibitor treatments. We treated WCRC-25, MDA-MB-134 and BT474 (positive control) cell lines with a PI3K inhibitor (Alpelisib) and an AKT inhibitor (AZD5363). As anticipated based on the RNA-sequencing data, WCRC-25 was sensitive to both inhibitors (Figure 6). WCRC-25 and MDA-MB-134 showed increased sensitivity in ULA compared to the 2D condition, perhaps due to the signaling supporting the anchorage-independent phenotype in these cell lines. Together, these results establish AKT pathway activation as a feature of metastatic progression and functionally validate PI3K/AKT inhibition as a candidate therapeutic strategy for WCRC-25.

**Figure 6.**
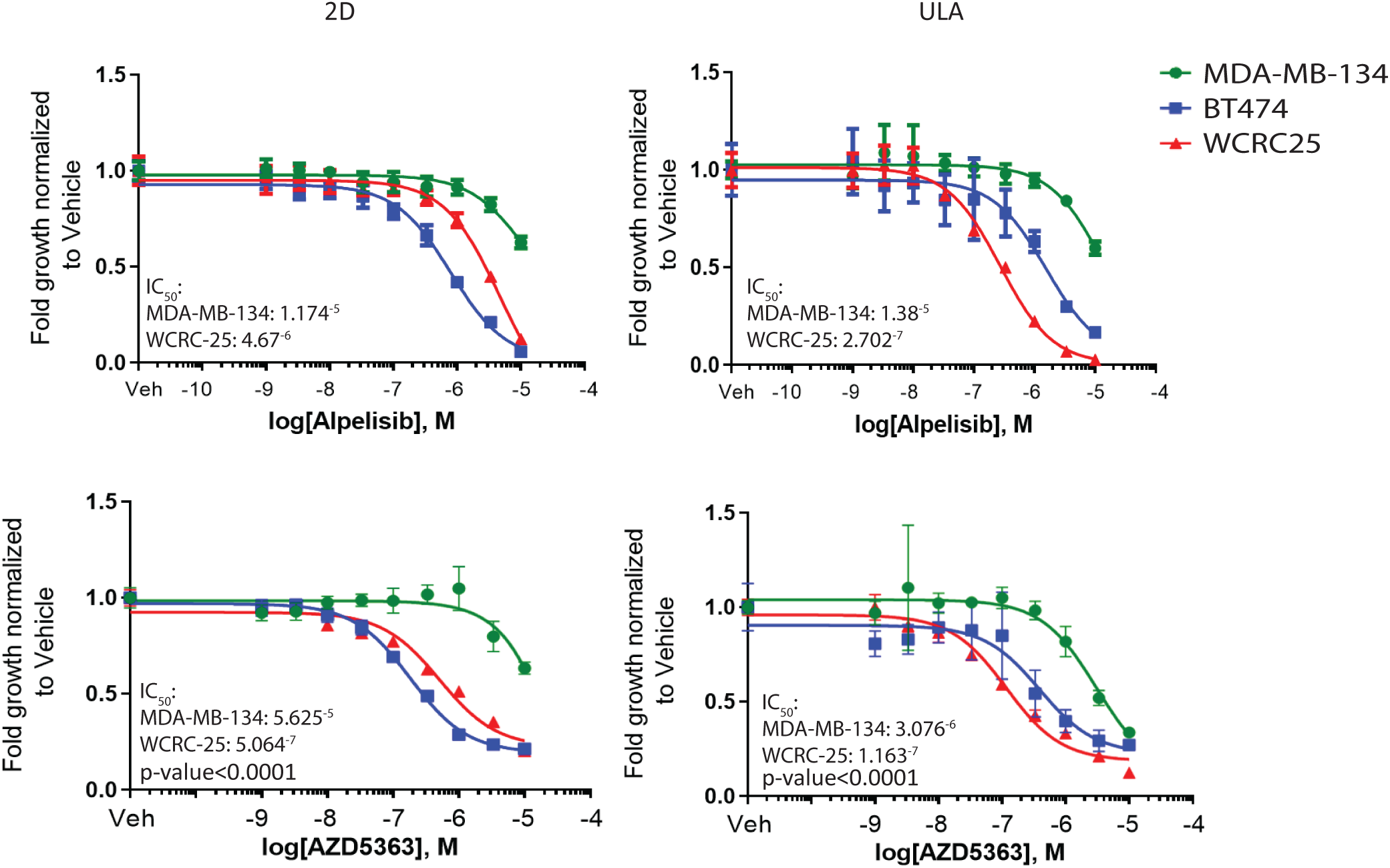
WCRC-25 shows susceptibility to PI3K and AKT inhibitors. (A) WCRC-25, MM134 and BT474 cells were seeded in 96-well 2D and ULA plates and treated with PI3K inhibitor (Alpelisib) or Akt inhibitor (AZD5363) for 6 days. CellTiter Glo assay was used to assess cell viability (relative luminescence) and data was normalized to vehicle treated control. IC50 values for viability were calculated by nonlinear regression and statistical differences between WCRC-25 and MM134 evaluated using sum-of-squares global F-test (p < 0.0001; representative experiment shown; n=3 (each with six biological replicates)).

### WCRC-25 grows and metastasizes in vivo

We assessed the ability of WCRC-25 to colonize and metastasize in vivo. We utilized traditional murine mammary fat pad (MFP) injection. In the MFP model, we had a 100% tumor take rate and rapid tumor growth (Figure 7A). There was no change in growth rates with the addition of estradiol (E2), as expected based on the lack of ER expression and response to ER antagonists in vitro (Figure 3A-B, Supplemental Figure S5A-B). The tumors displayed variable histologic patterns, with some featuring spindled cell metaplastic-like growth, some showing sheets of discohesive cells, and some showing the dispersed single-cell infiltration and single-file linear growth classically associated with lobular carcinoma (53) (Figure 7B) (additional examples are shown in Suppl Figure S7A). We also observed diverse histologies even within some individual tumors (Figure 7B, middle panels), as well as tumors with variable regions of abundant mucin production (Figure 7B, bottom panels). Many of the cells displayed mild to moderate anisocytosis and anisokaryosis, indistinct cell borders, and scant to moderate eosinophilic cytoplasm. Furthermore, we found that the WCRC-25 model spontaneously metastasized from the mammary gland to the lung in 3 of 9 mice (Figure 7C, and Suppl Figure S7B). No metastases were found in other organs such as the liver and ovaries. Overall, these data suggest that the WCRC-25 model recapitulates important aspects of the histology of ILC and may also be a tool for future spontaneous metastasis studies of ILC.

**Figure 7.**
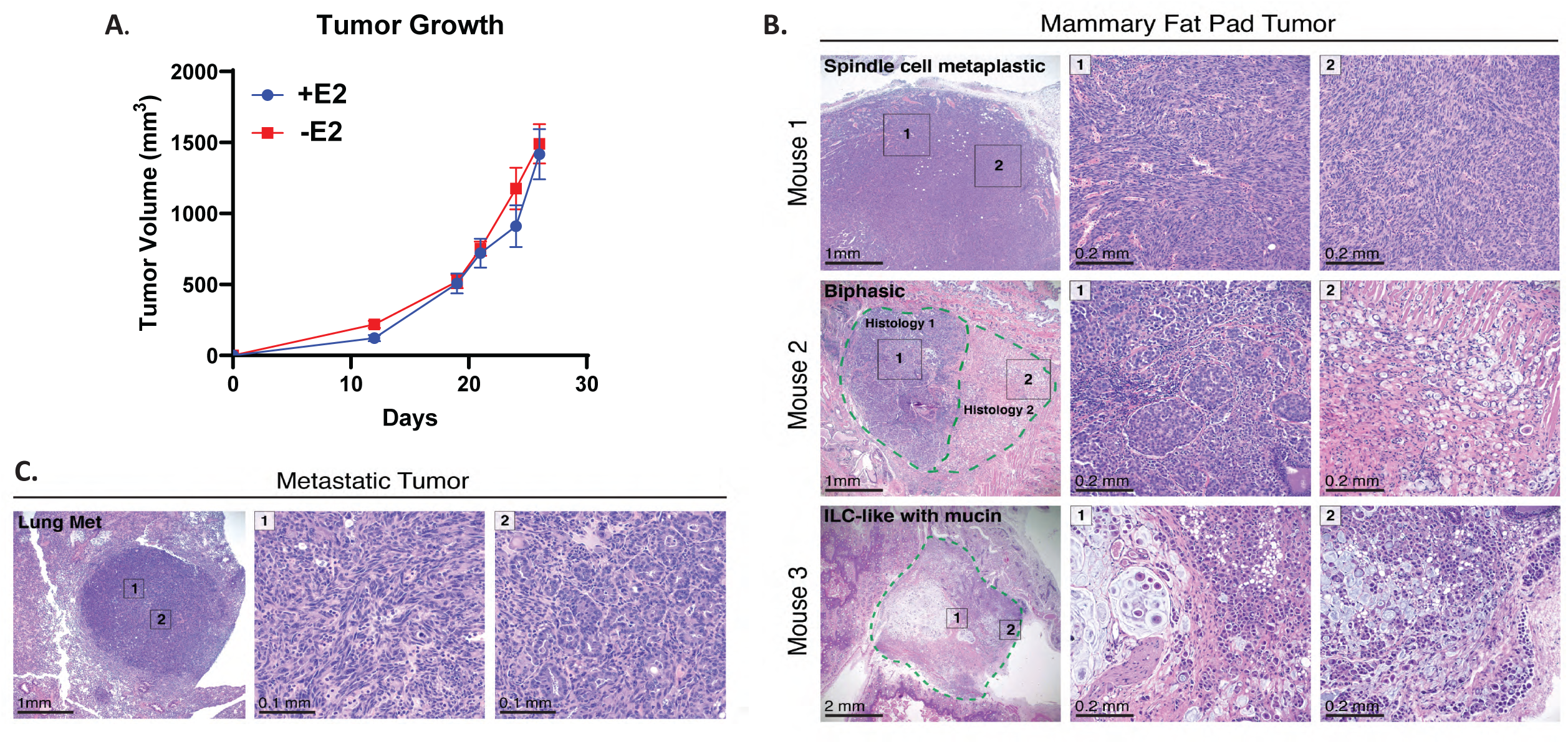
WCRC-25 cells grow rapidly *in vivo* and metastasize to the Lung. (A) Tumor volume over time with or without estradiol supplementation after mammary fat pad injection of WCRC-25 cells. (B) H&E stain of the primary tumors at endpoint. (C) H&E stain of a metastatic lesion within the lung.

In addition to the MFP model, we utilized the mouse intraductal (MIND) model (54). Prior to intraductal injections, WCRC-25 cells were labeled with GFP-Luc2 to track tumor progression *in vivo*. After 7 days, one mouse was sacrificed to determine disease progression at an early timepoint. We were able to detect GFP signal lining the mammary ducts (Figure 8A), and cancer cells within the ducts (Figure 8B), indicative of successful injection of WCRC-25 cells into the mammary ducts. Masson’s Trichrome staining revealed periductal collagen deposition (Figure 8C), which is in line with the extracellular matrix remodeling commonly described to occur in ILC (55).

**Figure 8.**
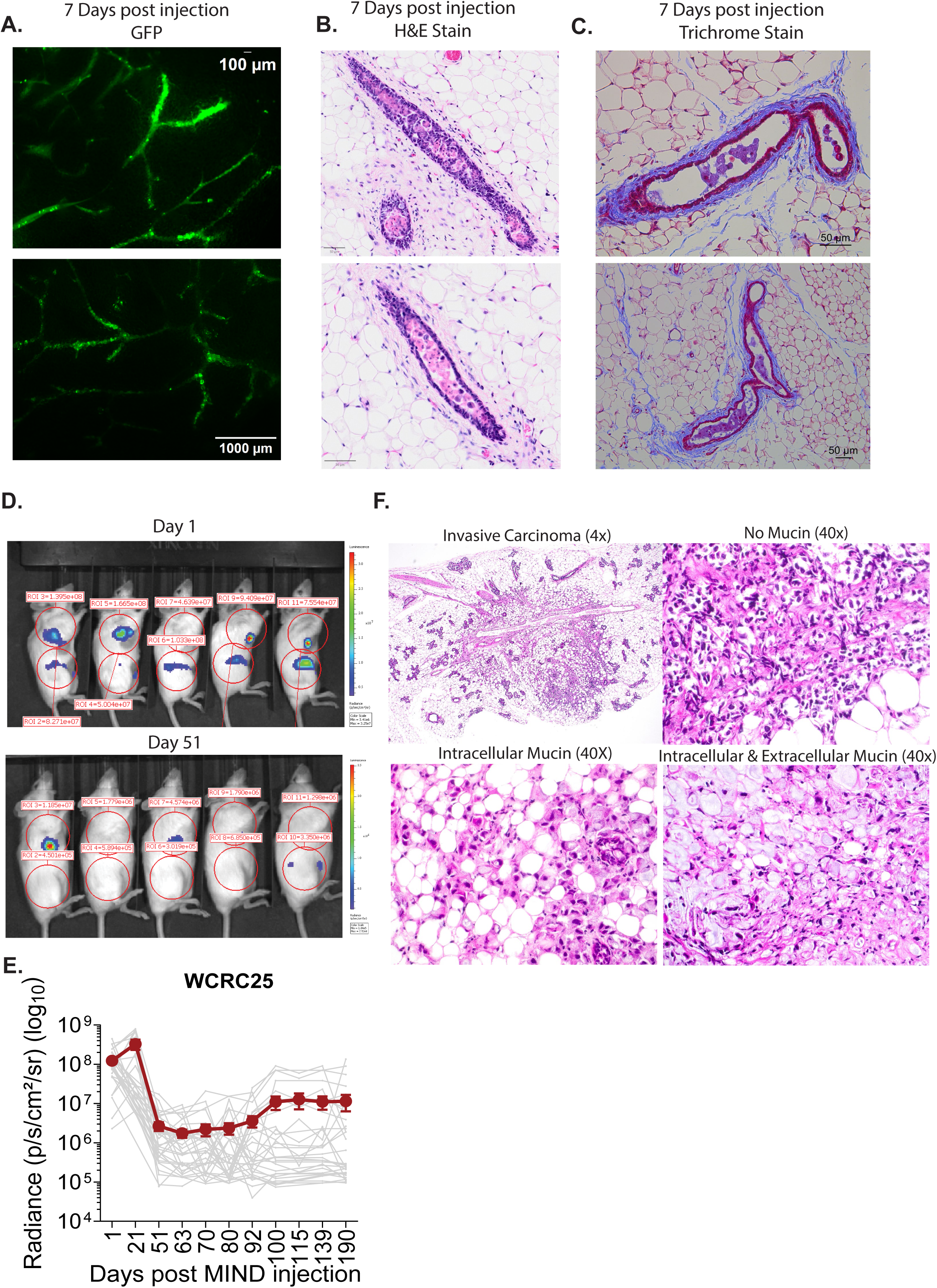
Engraftment of the WCRC-25 cells into the ducts of the mammary gland. (A) Immunofluorescent pictures of the GFP-Luc2 tagged WCRC-25 cells in the mammary gland 7 days after intraductal injections. (B) H&E staining of the mammary ducts 7 days after intraductal injection of WCRC-25 cells. (C) Trichrome staining of the mammary ducts 7 days after intraductal injection of WCRC-25 cells. (D) IVIS imaging of luciferase signal of mice injected with GFP-Luc2 tagged WCRC-25 cells 1 day and 51 days post intraductal injections. (E) Average luciferase signal (radiance) over time. (F) Analysis of tumors at 190 days. Top left: Representative H&E image of invasive lobular carcinoma (4x magnification). Top right: Representative H&E image of invasive lobular carcinoma with no mucin expression (40x magnification). Bottom left: Representative H&E image of invasive lobular carcinoma with intracellular mucin expression (40x magnification). Bottom right: Representative H&E image of the invasive lobular carcinoma with intracellular and extracellular mucin expression (40x magnification).

We had a 78% tumor take rate in the WCRC-25 MIND model. The luciferase signal peaked at 21 days, then regressed significantly, remained stable for ∼4 weeks, and subsequently began to increase again (Figure 8D-E). This is not a typical growth pattern seen in the MIND model (55), nor did we observe this regression in the mammary fat pad engraftments (Figure 7A). After ∼9 months, we harvested the largest remaining tumor and re-transplanted tumor slices into the fat pad of NSG mice to select for a more luminal-like phenotype.

However, we were unable to grow any tumors using this approach.

The remaining mammary fat pads were harvested and analyzed histologically for WCRC-25 cell colonization of the ducts and surrounding stroma. Three of nine mammary fat pads contained invasive carcinomas with lobular-like phenotypes infiltrating the adipose tissue. We observed a wide range of mucin production and cells with signet ring cytology. One invasive carcinoma had an abundance of mucin intracellularly and extracellularly, another invasive carcinoma had mucin that remained intracellular, and the third carcinoma had no detectable mucin (Figure 8F).

Together, the mammary fat pad and MIND models demonstrate that WCRC-25 recapitulates *in vivo* features of ILC, including ability to metastasize and lobular-type histology with variable mucin production, while also revealing model-specific growth dynamics that merit further investigation.

## Discussion

Cell lines remain a cornerstone of cancer research owing to their ease of use, low cost, and capacity for wide sharing, but only a handful of ILC cell lines are currently available (19). Here we report the generation and characterization of WCRC-25, a novel ER-negative ILC cell line with LumB/Her2 molecular subtype features. WCRC-25 was initially established from a pleural effusion using a modified Schlegel method but grew equally well in standard DMEM/10% FBS once established, indicating that continued use of CM-ROCKi conditions is unnecessary after outgrowth. WCRC-25 also expressed the basal marker CK14, which initially seemed unexpected but is consistent with reports that a subset of ILC tumors retain basal cytokeratin expression - 17% of cases in one 82-tumor series (56) - although another study reported absence of CK5/6, CK14 and CK17 in ILC (57), and basal marker expression in luminal tumors has even been linked to improved survival (58). These discrepancies may partly reflect that our sample, unlike most published series, derives from metastatic rather than primary ILC.

Doubling time, morphology, and anchorage-independent growth in WCRC-25 were comparable to existing ILC models (24, 59). Its significant migration toward Collagen I is notable, as only two of the four currently available ER+ ILC cell lines show this capability (24); the ER-negative lines UACC-3133 and MA-11 have not yet been evaluated for this phenotype and warrant future study.

Longitudinal DNA panel and RNA sequencing of patient specimens tracked mutations, receptor tyrosine kinase signaling, and cell-cycle signatures through disease progression, while optical genome mapping of WCRC-25 revealed structural variants not previously described. Notably, the *CDH1* Q706* mutation identified in WCRC-25 was also detectable in the patient’s cell-free DNA, supporting the feasibility of liquid biopsy for tracking ILC progression (60, 61). A limitation of our study is that, apart from *CDH1* (confirmed by Sanger sequencing and ddPCR), other genetic alterations were not orthogonally validated; differences in sequencing technology and filtering pipelines can introduce false positive or negative calls.

Guided by our omics data, we identified PI3K/AKT signaling as a candidate therapeutic vulnerability; WCRC-25 showed sensitivity to PI3K and AKT inhibitors comparable to BT474, a *PIK3CA*-mutant control line. Several other findings from our mutational and transcriptomic analyses merit further mechanistic follow-up, including the Her2 features of the cell line.

WCRC-25 engrafted efficiently in both the mammary fat pad and the mammary ducts, forming tumors with histologic features of ILC. Mammary fat pad engraftment produced rapid tumor growth and pulmonary metastases in one-third of mice within approximately three weeks. Additional studies with earlier tumor removal and longer observation are required to capture additional metastases, including to sites more commonly affected in ILC patients such as bone and reproductive tract, testing the utility of this model for studying ILC metastasis. Intraductal (MIND) engraftment was initially successful but showed an atypical luciferase signal that regressed after three weeks before increasing again, unlike the steady growth typical of NST and ILC lines in this model (55); this pattern may reflect intrinsic WCRC-25 biology, an immune response to the GFP-luciferase tag, or both, and an untagged control line would help distinguish these possibilities. It is worth noting that a basal differentiation program has previously been associated with growth in the fat pad and inhibited intraductal growth(54). Given the expression of basal cytokeratins in WCRC-25, this program could also play a role in this model.

MIND tumors also displayed a range of mucin production and signet-ring morphology, including one tumor with abundant extracellular mucin, a feature only rarely reported in ILC (62). Together with recently described ILC xenograft models (63–65), WCRC-25 expands the translational toolkit available for *in vivo* ILC research.

In conclusion, WCRC-25 is a novel ER-negative, LumB/Her2 ILC cell line that we have characterized phenotypically and genomically, and that we believe will be a valuable resource for future studies of ILC biology and its therapeutic vulnerabilities such as drugs targeting growth factor receptor pathways.

## Authors’ Contributions

Conception and design: E.A. Bossart, N. Tasdemir, S.L. Puhalla, J.M. Atkinson, A.V. Lee, S. Oesterreich

Development of methodology: E.A. Bossart, N. Tasdemir, T. Heim, L. Miller, S.L. Puhalla, J.M. Atkinson, A.V. Lee, S. Oesterreich

Acquisition of data: A. Elangovan, E.A. Bossart, N. Tasdemir, A. Basudan, O.S. Shah, C. Meier, T. Liu, L. Brown, S. Attaran, K. Ding, C. Meier, G Sflomos, L. Miller, K Kota, T. Liu, P.C. Lucas, P.F. McAuliffe

Analysis and interpretation of data: A. Elangovan, E.A. Bossart, N. Tasdemir, A. Basudan, C. Neumann, O.S. Shah, T. Liu, S.C. Purdy, K. Ding, C. Meier, L. Rigatti, G Sflomos, C Brisken, T. Liu, G. Gurda

Writing, review, and/or revision of the manuscript: A. Elangovan, E.A. Bossart, O.S. Shah, S.C. Purdy, J. Hooda, GT Gurda, K. Ding, J.M. Atkinson, A.V. Lee, S. Oesterreich

Administrative, technical, or material support: N. Tasdemir, C. Meier, T. Heim, L. Brown, J. Hooda, L. Miller, S.L. Puhalla, G. Gurda, P.F. McAuliffe

Study supervision: E.A. Bossart, N. Tasdemir, S.L. Puhalla, P.F. McAuliffe, J.M. Atkinson, A.V. Lee, S. Oesterreich.

## Acknowledgements

We are forever grateful to the patient for donating her tissue for research. We are also thankful to Jian Chen, Julie Scott, and Weizhou Hou for outstanding technical support. Thanks to the University of Pittsburgh Genomics Research Core (GRC) for assistance with MammaSeq-based DNA sequencing and Illumina SNP array analysis, and to the UPMC Hillman Cancer Center and Tissue and Research Pathology/Pitt Biospecimen/Animal Facility Cores shared resource, which are supported in part by award P30CA047904. We would like to also thank Bionano Genomics for collaboration and assistance with optical genome mapping. This work was supported in part by a Susan G Komen Leadership Grant to SO (SAC160073) and NCI R01 CA252378 (SO/AVL). AVL and SO are Komen Scholars, Hillman Foundation Fellows, and are supported by BCRF. This research was also supported in part by the University of Pittsburgh Center for Research Computing (RRID:SCR_022735), including use of the HTC cluster, which is supported by NIH award number S10OD028483. Partial support was also provided by the Magee-Womens Research Institute and Foundation, the Shear Family Foundation, and the Metastatic Breast Cancer Network.

---

**Supplemental Figure 1.**
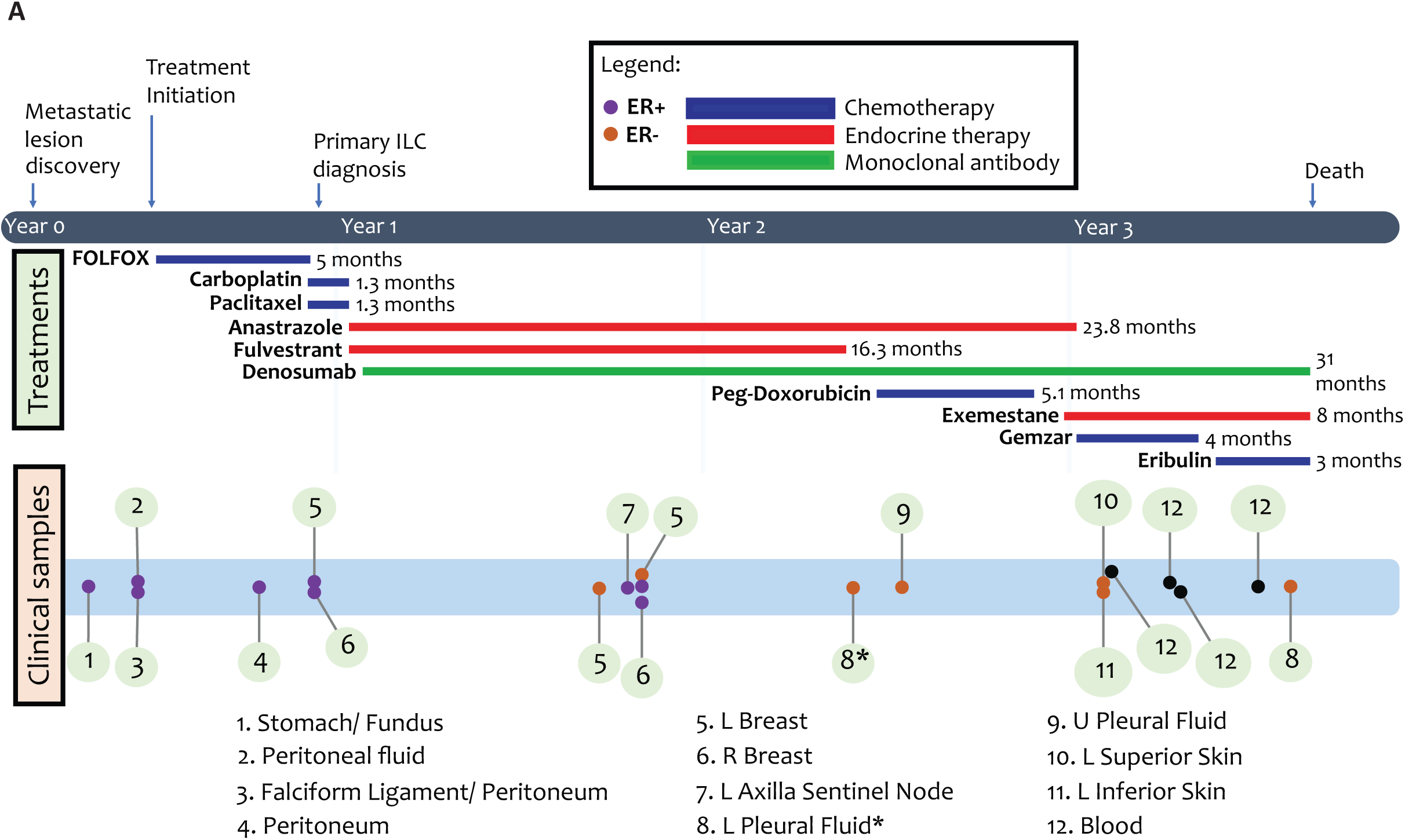
A schematic representation of WCRC-25 patient clinical history and longitudinal sample collection timeline with the respective analyses performed annotated. The “*” indicates the source of the cell line.

**Supplemental Figure 2.**
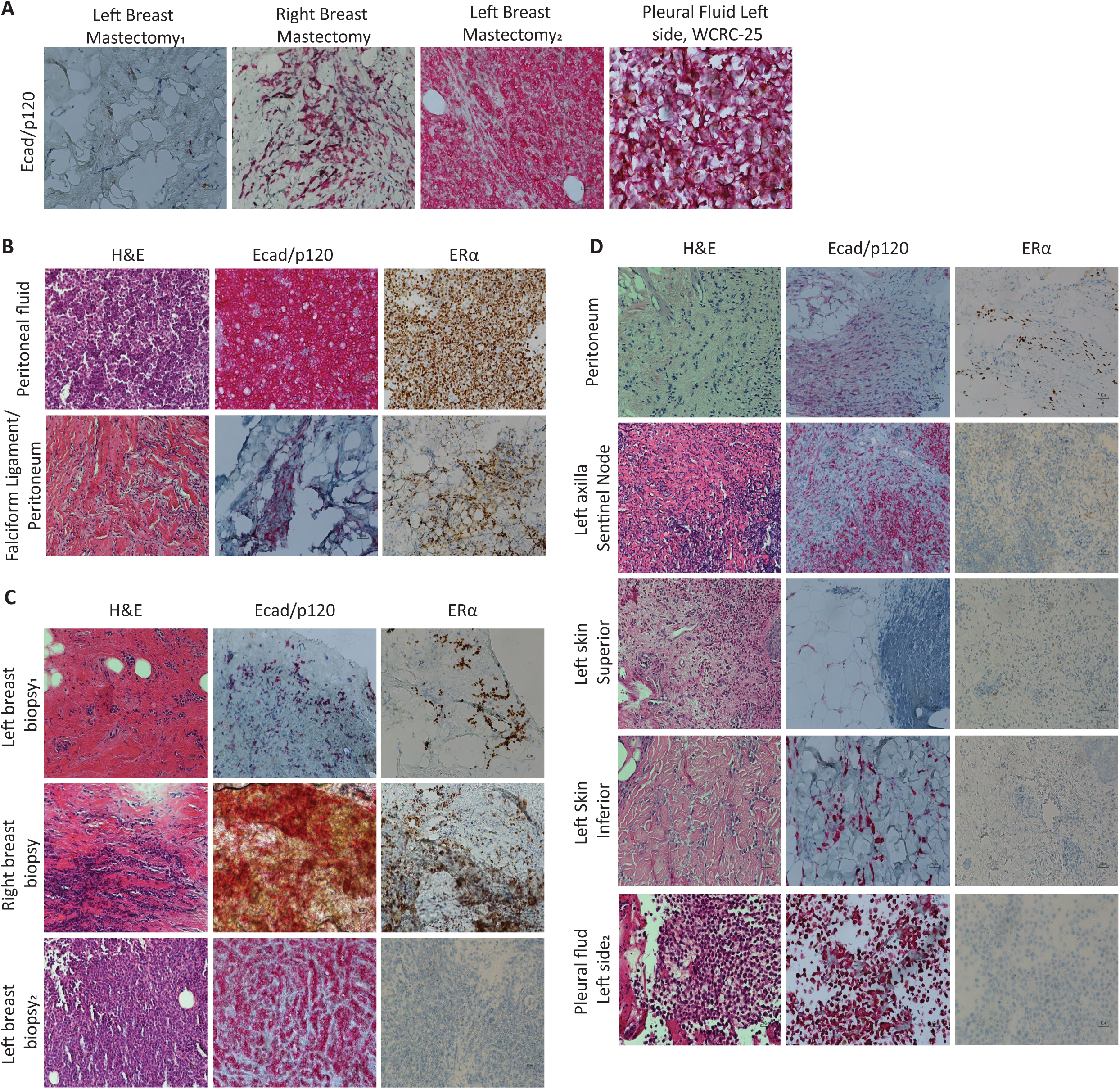
H&E and E-cad/p120 and ERα IHC staining of (A) primary tumors and PE, (B) therapy naïve samples, (C) breast biopsy samples and (D) chemotherapy or endocrine therapy treated metastatic lesions from the patient (20X magnification; bars represent 40 µm). Of note, the quality of remaining tissues used for some of the staining was limited e.g left mastectomy^1^ but it was included since even in limited cancer cells the loss of E-cad and translocation of p120 was visible.

**Supplemental Figure 3.**
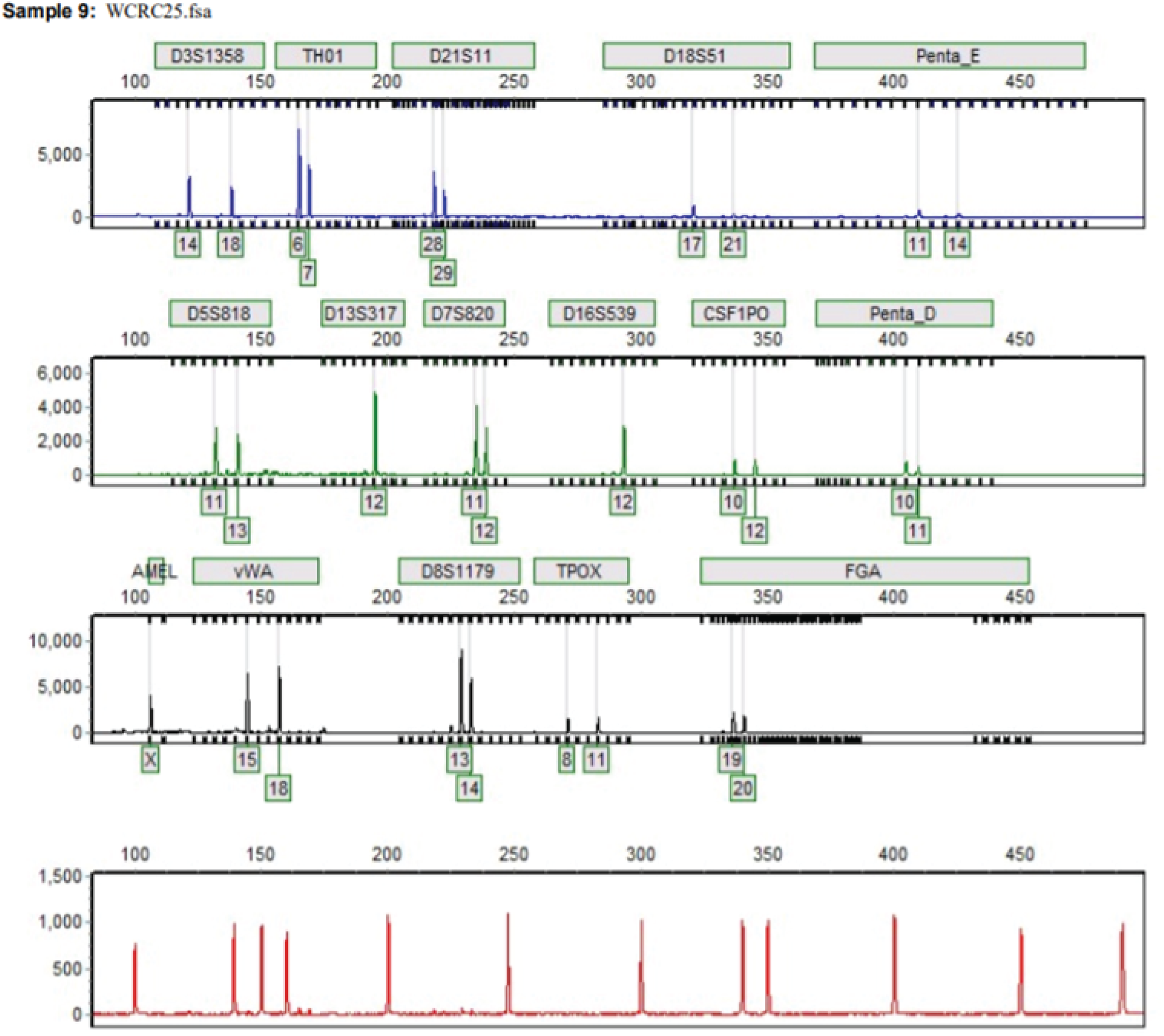
STR analysis of WCRC-25 shows the novelty of the cell line.

**Supplemental Figure 4.**
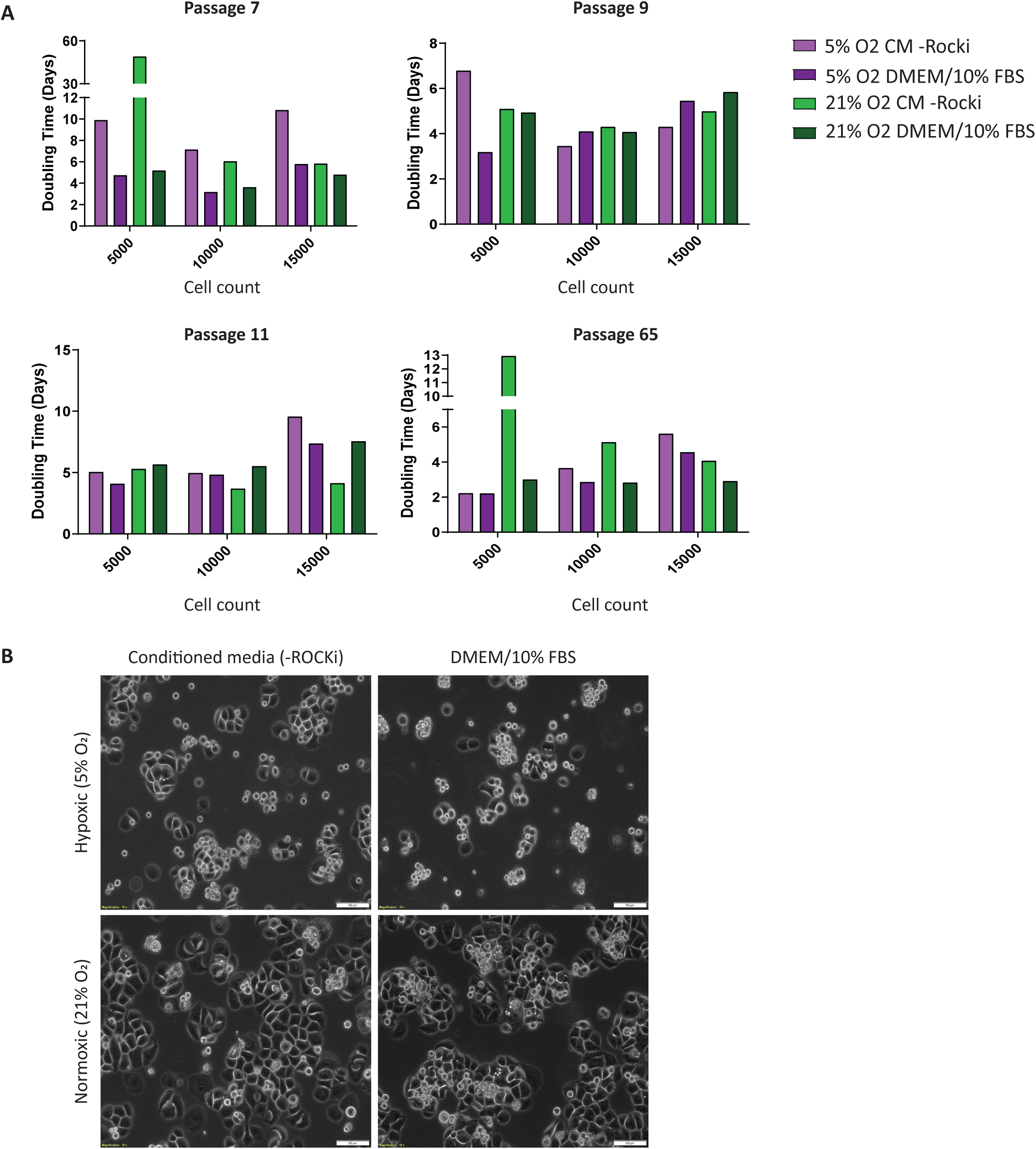
(A) Cell proliferation assay performed with WCRC-25 cells at passages 7, 9, 11 and 65 post generation to assess cell doubling times. Quantification was performed with FluoReporter™ Blue Fluorometric dsDNA Quantitation Kit (Invitrogen #F2962). Several culture conditions with differing O2, CO2, media and ROCK inhibitor presence were assessed, imaged in (B).

**Supplemental Figure 5.**
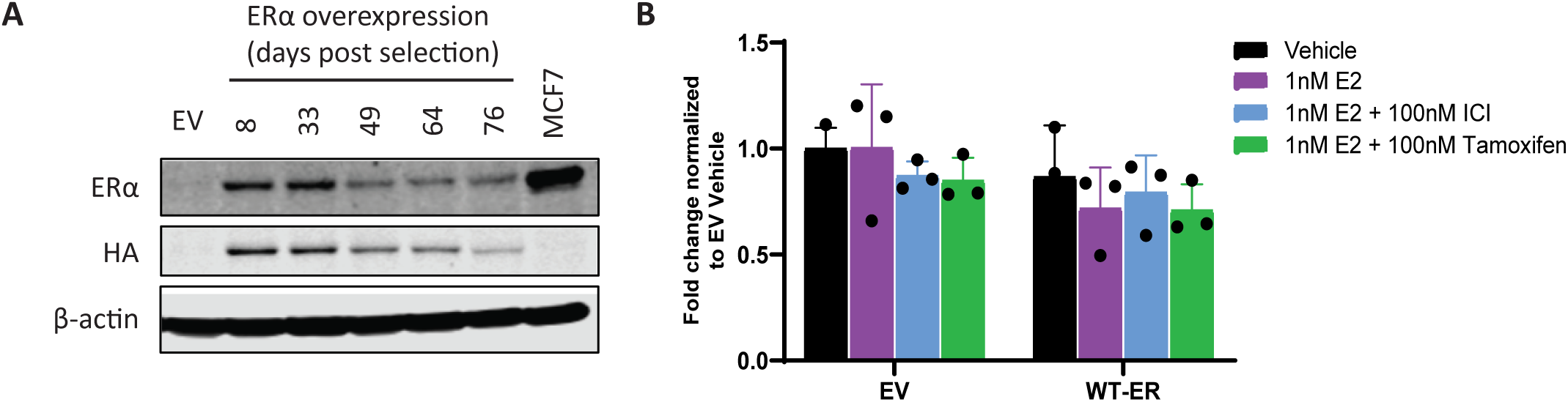
(A) Immunoblot for ERα, HA and β-actin showing the loss of ERα overexpression levels over time. MCF7 cell lysate was used as a positive control for ERα. (B) WCRC-25 cell line +/-ERα overexpression was subjected to an ERE-TK assay after 3 days of hormone deprivation followed by 1 nM E2, 100 nM ICI or 100 nM Tamoxifen treatments for 24 hours.

**Supplemental Figure 6.**
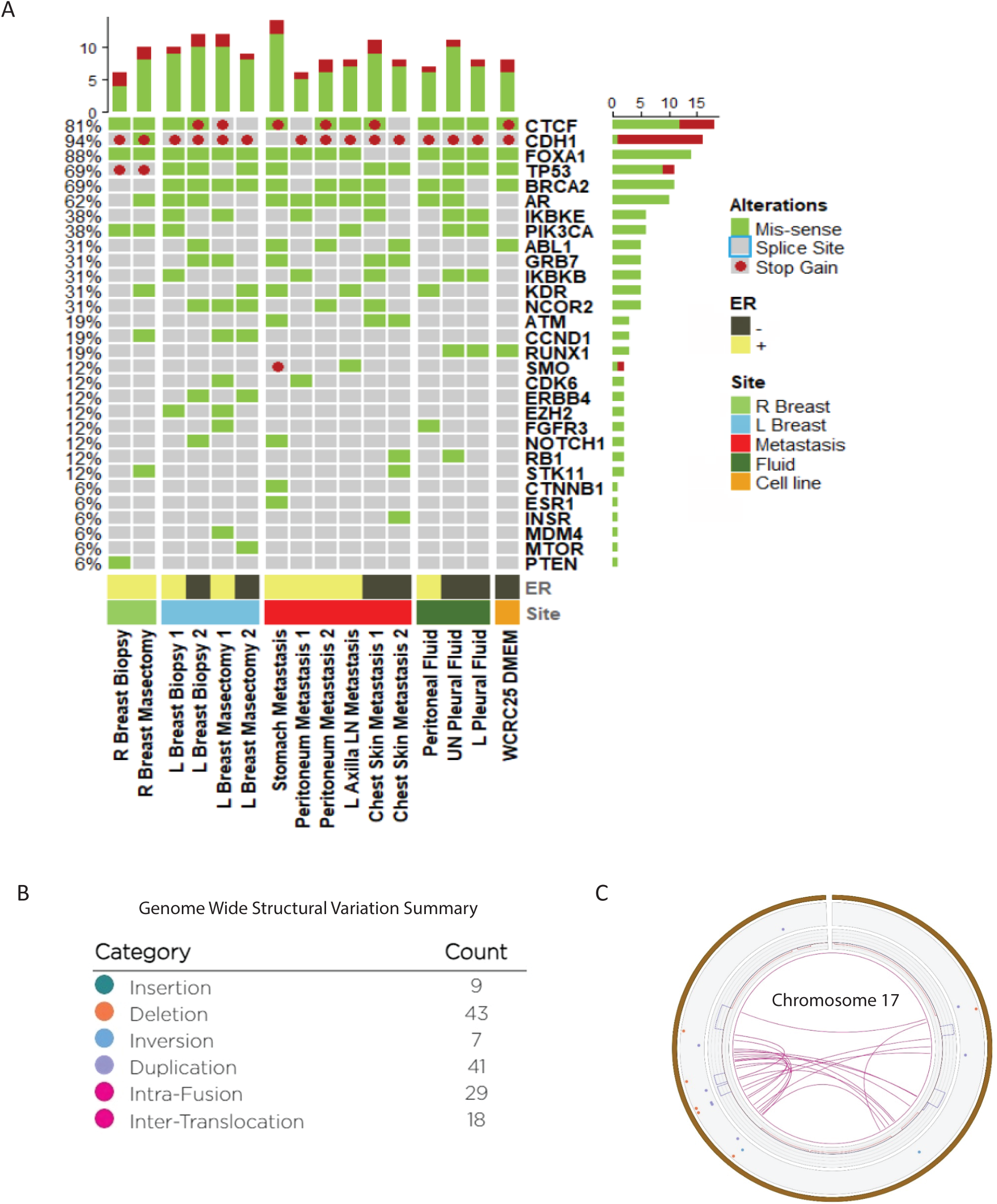
(A) Oncoprint showing top 30 alterations identified using MammaSeq. Each row shows a unique mutation (labelled with gene name and along with the mutation induced protein change). Each column represents a sample. Samples are ordered as those from right breast, left breast, metastasis, fluid and cell line. (B) Summary of all SV types and their count is shown in table in bottom right. (C) Circos plot of intra-chromosomal translocations in chromosome 17 of WCRC-25 cell line.

**Supplemental Figure 7.**
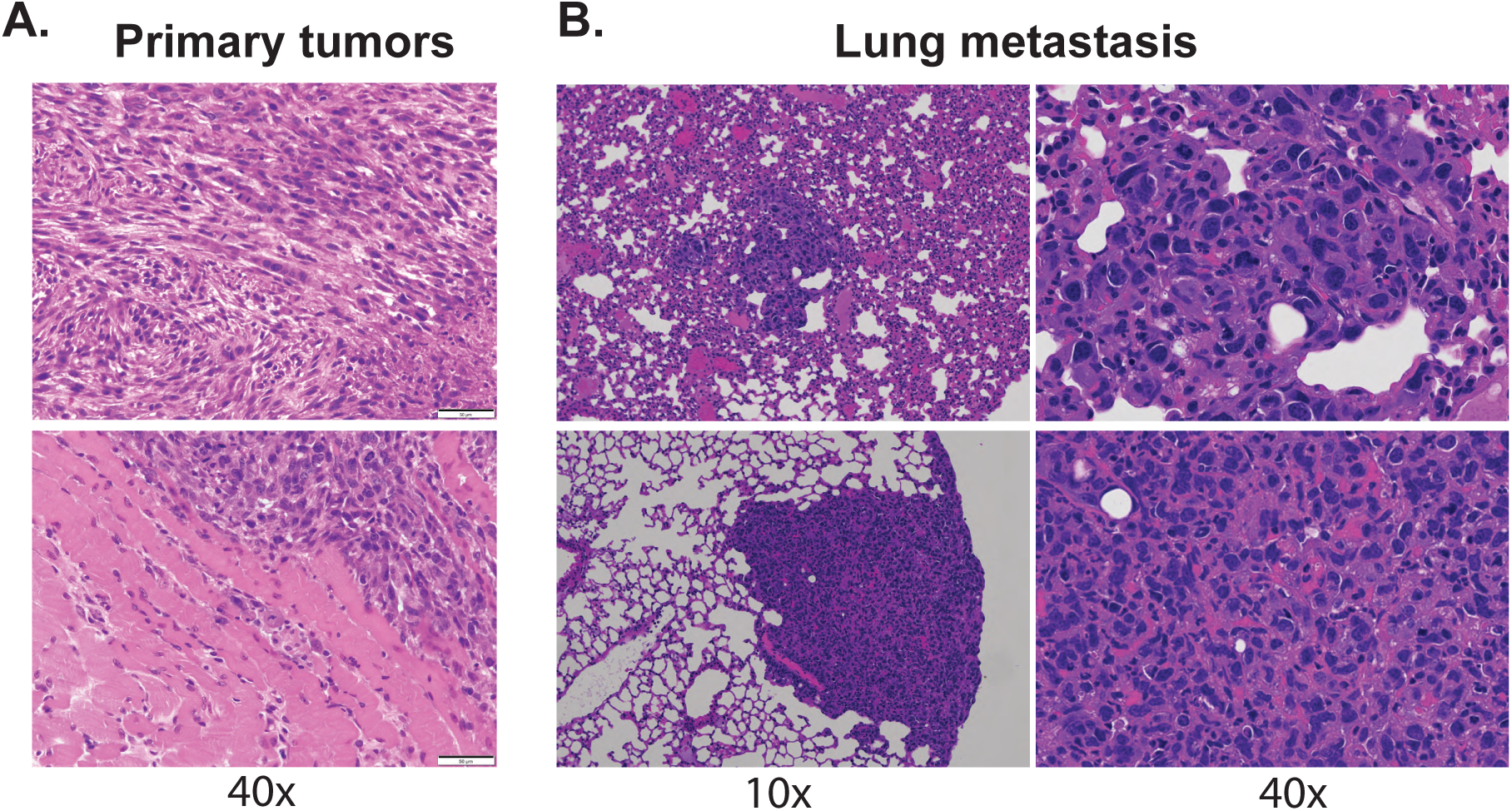
(A) H&E stain of the primary tumors at endpoint (40x magnification). (C) H&E stain of the metastatic lesions within the lung (10x and 40x magnification).

